# Multivalent Exosome based protein vaccine: a “mix and match” approach to epidemic viruses’ challenges

**DOI:** 10.1101/2023.07.13.548895

**Authors:** Mafalda Cacciottolo, Li-En Hsieh, Yujia Li, Michael J. LeClaire, Justin B Nice, Ryan Twaddle, Ciana Mora, Meredith Young, Jenna Angeles, Christy Lau, Cindy Jin, Kristi Elliott, Minghao Sun

## Abstract

Endemic viruses are becoming increasingly the norm, and the development of a rapid and effective vaccine is emergent. Here, we used our StealthX ™ exosome platform to express either Influenza H3 (Stealth™ X-Hemagglutinin, STX-H3) or SARS-CoV-2 Delta spike (Stealth™ X-Spike, STX-S) protein on the surface and facilitate their trafficking to the exosomes. When administered as single product, both STX-H3 and STX-S induced a strong immunization with the production of a potent humoral and cellular immune response in mice. Interestingly, these effects were obtained with administration of nanograms of protein and without adjuvant. Therefore, we tested the possibility of a multivalent vaccine: STX-H3 and STX-S exosomes were formulated together in a “mix and match” approach and the immune response was further evaluated. We showed that our STX-H3+S cocktail vaccine is as effective as the single components administered separately, resulting in a strong antibody and T-cell response. Our data show that our exosome platform has an enormous potential to revolutionize vaccinology by rapidly facilitating antigen presentation, and for therapeutics by enabling cell and tissue specific targeting.

## INTRODUCTION

In the aftermath of the initial emergence of SARS-CoV-2 in November 2019, it is evident that the virus is becoming endemic, a status already held by Influenza and respiratory syncytial virus. Similar to the Influenza A/H3N2 viruses, the most rapid changing subtype, more than a dozen main variants that have been reported for SARS-CoV-2 virus (cdc.org), with increasing number of breakthrough infections and consequent implication on the efficacy of the vaccines. An example is offered by the recent BQ.1 and XBB variants: both were found to be highly resistant to the polyclonal sera and the monoclonal antibodies that had appreciable neutralizing activities against the initial Omicron variant [1, 2]. With the disease becoming endemic, a vaccine that can be adapted quickly alongside the evolution of both SARS-CoV-2 and Influenza is extremely important.

Before and after the COVID-19 pandemic, influenza was one of the most important and frequent viral disease of the respiratory system. All persons are recommended to receive the influenza vaccine, especially children ≤ 5 years old, adults ≥ 50 years old, patients with chronic conditions or who are immunocompromised, and women who are or will be pregnant during the influenza season [3]. Due to the characteristics of influenza virus replication, mutation of viral genome is frequent [4], and vaccination against new strains is needed annually especially in the cohorts with higher risk of severe diseases [3]. The prediction for the vaccine strain is based on the epidemiological monitoring of the circulating strains and the vaccine strains need to be updated frequently [3]. Therefore, it is necessary for influenza vaccines to be able to quickly adapt to combat the ever-changing virus. Moreover, a bivalent vaccine would be highly preferred to reduce numbers of injections and costs, wherever efficacy against the antigens is preserved.

We have previously showed that our exosome platform StealthX™ can be used to generate a novel protein-based vaccine able to elicit a potent cellular and humoral immune response against SARS-CoV-2 spike and nucleocapsid [5, 6]: both antigens were combined together with no effect on the immune response to each other, suggesting no immune competition between antigens from the same virus.

In the present study, we sought to use our exosome platform to generate immune response against both SARS-CoV-2 and influenza virus. Our results showed that mice immunized with exosomes carrying SARS-CoV-2 spike protein or influenza A (H3N2) hemagglutinin elicited potent cellular immune response and produced high titer of antibodies against target immunogens. Furthermore, mice that received the bivalent formulation also showed a synergistic cellular and humoral immune response toward SARS-CoV-2 spike and influenza A hemagglutinin. Our data showed that the exosome-based STX vaccine platform is able to elicit robust humoral and cellular immune response against different viral antigen simultaneously with low antigen doses in the *in vivo* mouse model. The platform can be potentially adapted to different target antigens or mutants to combat the fast-evolving viral infections.

## METHODS

### Cell lines

Human embryonic kidney 293 T cells (293T) were purchased from ATCC (CRL-3216). 293T cells were maintained in culture using Dulbecco’s Modified Eagle Medium (DMEM), high glucose, Glutamax™ containing 10% fetal bovine serum. 293T cells were incubated at 37°C /5% CO_2_. FreeStyle™ 293F cells (Gibco, 51-0029) were purchased from ThermoFisher. 293F cells were used as a parental cell line to generate spike SARS-CoV-2 delta spike and Influenza A virus hemagglutinin expressing stable cell lines: Stealth X-Spike cells (STX-S) and Stealth X-H3 cells (STX-H3). 293F, STX-S, and STX-H3 cells were maintained in a Multitron incubator (Infors HT) at 37°C, 80% humidified atmosphere with 8% CO_2_ on an orbital shaker platform rotating at 110 rpm.

### Lentiviral vectors

Lentiviral vectors for expression of SARS-CoV-2 spike (Delta variant B.1.617.2, NCBI accession # OX014251.1) and Influenza virus hemagglutinin (A/New Yotk/392/2004, NCBI accession # YP_308839) were designed and synthesized from Genscript together with the two packaging plasmids (pMD2.G and psPAX2). Lentiviral particles for transduction were generated as previously (Cacciottolo et al, under revision). Briefly, lentiviral particles for transduction were generated by transfecting 293T cells with pMG.2 (Genescript), psPAX2 (Genescript) and STX-S_pLenti (Genscript) expressing spike or STX-H3_pGenlenti (Genscript) at a ratio of 5:5:1 using Lipofectamine 3000 according to the manufacture’s instruction. Spike and H3 lentiviral particles were collected at 72 hours post transfection and used to transduce 293F parental cells to generate STX-S and STX-H3 respectively.

### Flow cytometry

Standard flow cytometry methods were applied to measure the expression of SARS-CoV-2 spike protein and Influenza A hemagglutinin protein on STX-S and STX-H3 cell surface, respectively. In brief, 250K STX cells were aliquoted, pelleted and resuspended in 100μL eBioscience™ Flow Cytometry Staining Buffer (ThermoFisher). Cells were incubated at room temperature (RT) for 30 min protected from light in the presence of anti-spike antibody (Abcam, clone 1A9, ab273433) or anti-H3 (A/Perth/16/2009) (Immune technology, IT-003-004M18) labeled with Alexa Fluor®-647 (Alexa Fluor® 647 Conjugation Kit (Fast)-Lightning-Link® (Abcam, ab269823) according to the manufacturer’s protocol. Following incubation, STX cells were washed with eBioscience™ Flow Cytometry Staining Buffer (ThermoFisher, cat No 00-4222-57), resuspended in PBS and analyzed on the CytoFlex S flow cytometer (Beckman Coulter). Data was analyzed by FlowJo (Becton, Dickinson and Company; 2021).

### Cell sorting

Cell sorting was performed at the Flow Cytometry Facility at the Scripps Research Institute (San Diego, CA). To enrich the spike positive population, STX-S cells were stained as described above for flow cytometry and went through cell sorting (Beckman Coulter MoFlo Astrios EQ) to generate pooled STX-S. The pooled STX-S were used in all the experiments in this paper unless specified otherwise.

### STX exosome production

STX-S and STX-H3 cells were cultured in FreeStyle media (ThermoFisher, 12338018) in a Multitron incubator (Infors HT) at 37°C, 80% humidified atmosphere with 8% CO_2_ on an orbital shaker platform. Subsequently, cells and cell debris were removed by centrifugation, while microvesicles and other extracellular vesicles larger than ∼220 nm were removed by vacuum filtration. Next, exosomes were isolated using either Capricor’s lab scale or large-scale purification method. For lab scale: supernatant was subjected to concentrating filtration against a Centricon Plus-70 Centrifugal filter unit (Millipore, UFC710008), then subjected to size exclusion chromatography (SEC) using a qEV original SEC column (Izon, SP5). For large scale: supernatant was subjected to concentrating tangential flow filtration (TFF) on an AKTA Flux s instrument (Cytiva, United States) and then subjected to chromatography on an AKTA Avant 25 (Cytiva, United States).

### Nanoparticle tracking analysis

Exosome size distribution and concentration were determined using ZetaView Nanoparticle Tracking Analysis (Particle Metrix, Germany) according to manufacturer instructions. Exosome samples were diluted in 0.1 μm filtered 1X PBS (Gibco, 10010072) to fall within the instrument’s optimal operating range.

### Jess automated western blot

Detection of SARS-CoV-2 spike and H3 proteins in cell lysate and exosomes used Protein Simple’s Jess capillary protein detection system. Samples were lysed in RIPA buffer (ThermoFisher Scientific, 8990) supplemented with protease/phosphatase inhibitor (ThermoFisher Scientific, A32961), quantified using the BCA assay (ThermoFisher Scientific, 23227) and run for detection. To detect spike, the separation module 12–230 kDa was used following manufacturers protocol. Briefly, 0.8 μg of sample and protein standard were run in each capillary, probed with anti-mouse Ms-RD-SARS-COV-2 (R&D Systems, MAB105401, 1:10 dilution) or anti-mouse H3 (Immune-Tech, IT-003-004M2, 1:50 dilution) followed by secondary antibody provided in Jess kits (HRP substrate used neat).

### CD81 bead-assay

STX-S, STX-H3 or 293F parental exosomes were mixed with anti-CD81 labeled magnetic beads for 2 h at RT (ThermoFisher, 10622D) and washed twice with PBS using the magnetic stand. Next, the bead-exosomes were incubated with either direct conjugated AlexaFluor 647 anti-spike (see above flow cytometry), AlexaFluor 647 anti-H3, FITC anti-CD81 antibody (BD Biosciences, 551108), or FITC Mouse IgG, κ isotype control (BD Biosciences, 555748) for 1 h at RT followed by two PBS washes. 293F exosomes were used as a negative control for spike expression and the isotype antibody was used as a negative control for CD81 expression. Samples were analyzed on a CytoFlex S (Beckman Coulter) flow cytometer and data was analyzed by FlowJo.

### Animal studies – Mice

To examine the efficacy of STX exosomes, age matched BALB/c mice (female, 8-10 wks old) were anesthetized using isoflurane and received bilateral intramuscular injection (50 μl per leg, total 100 μl) of either 1) PBS 2) STX-S, 3) STX-H3 or 4) STX-H3+S exosomes. The booster injection was performed at day 21. Mice were monitored closely for changes in health and weight were recorded biweekly. Blood collection was performed at day 14 and day 35. Blood (∼50-500 μl) was collected from the submandibular vein and processed for plasma isolation after centrifugation at 4000 rpm for 5 min at 4°C. Timeline of mouse study is outlined in Fig. 2A.

**Figure 1.**
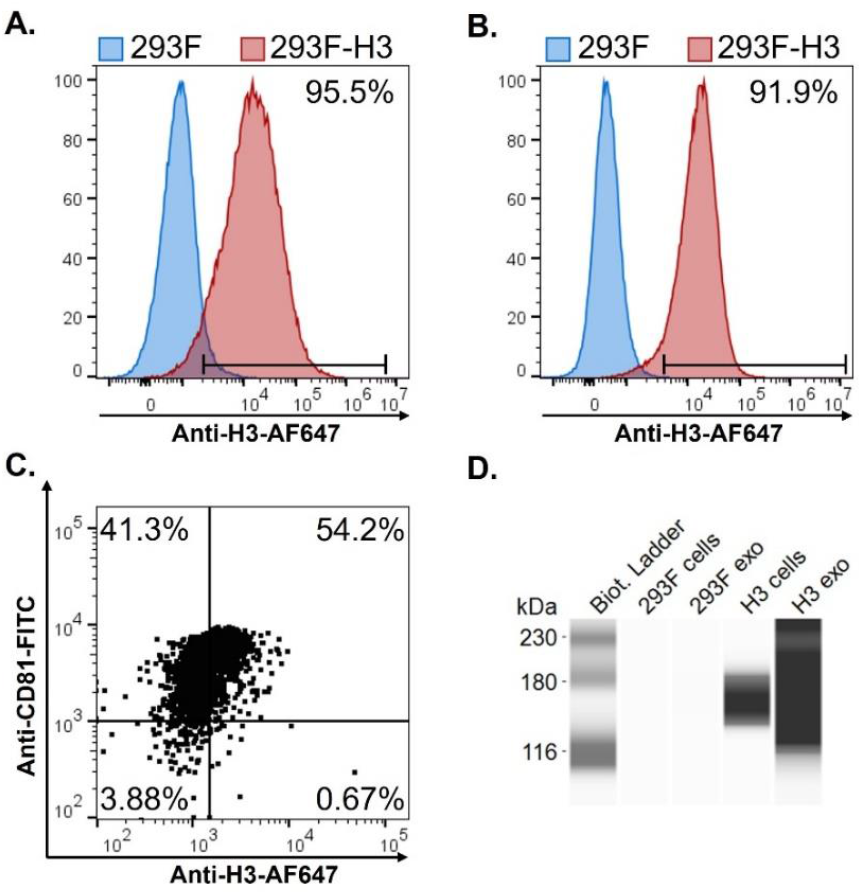
Characterization of STX-H3 cells and exosomes expressing hemagglutinin H3. **A**. High expression of H3 was detected on cell surface (red) by flow cytometry. Parent, non-engineered 293F cells (blue) do not express H3 protein, as expected. **B**. High expression of H3 was detected on exosome surface (red) by flow cytometry. Parent, non-engineered 293F exosome (blue) do not express H3 protein, as expected. **C**. Colocalization of CD81 and H3 protein on exosomes was confirmed by flow cytometry using a bead-assay. **D**. Enrichment of H3 protein in exosomes was confirmed by Jess-automated Western Blot. From left to right: Lane 1: marker, Lane 2: non-engineered 293F cells, Lane 3: non-engineered 293F exosomes, Lane 4: STX-H3 cells, Lane 5: STX-H3 exosomes.

**Figure 2.**
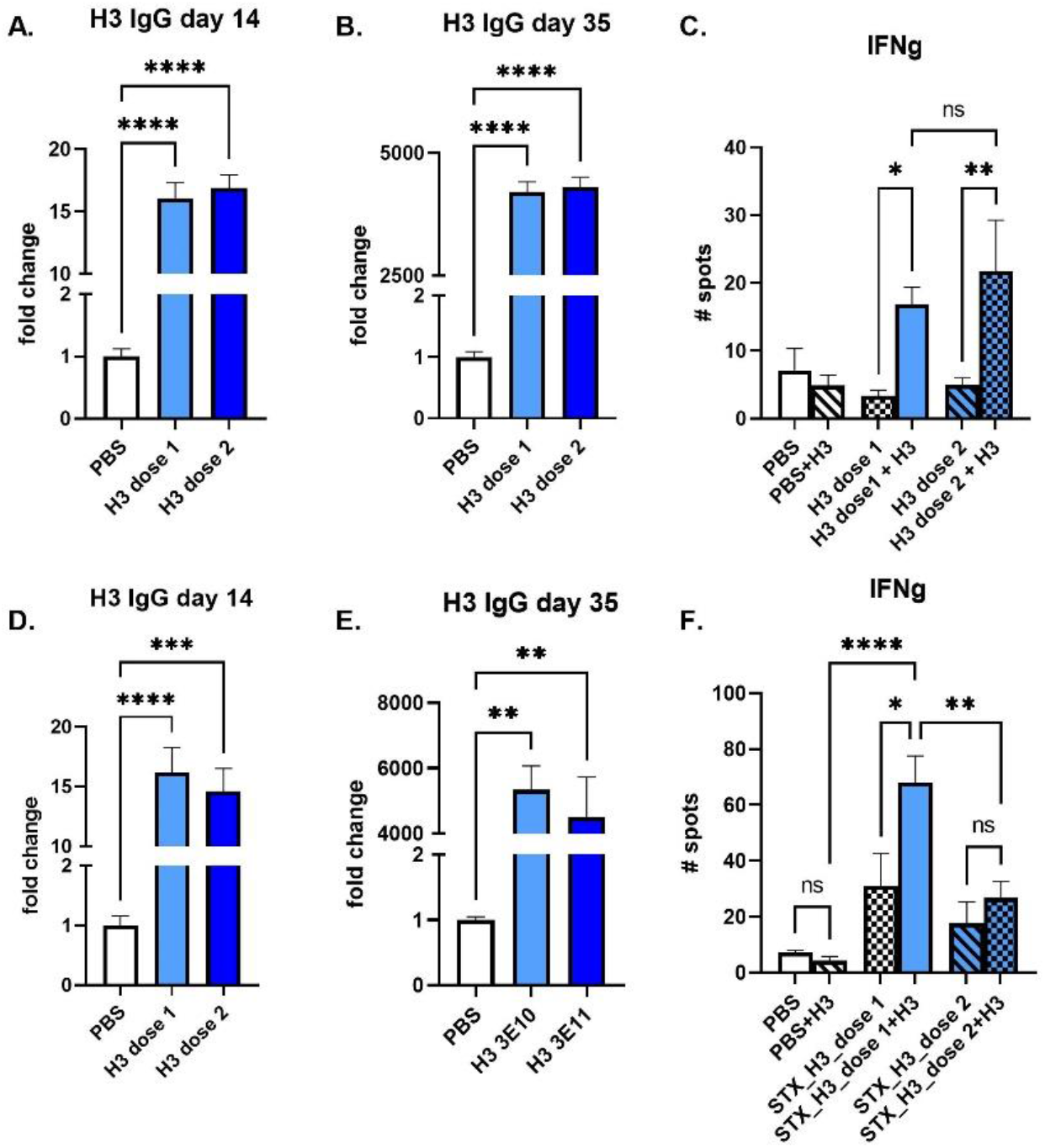
STX-H3 exosome vaccine elicited a robust immune response. The STX-H3 vaccine induced robust expression of anti H3 antibodies in mice after 1 (day 14) and 2 (day 35) IM injections as analyzed by ELISA in both young (A-B) and old mice (D-E). Additionally, a strong T-cell response is observed (C, D). PBS was used as a vehicle control in all studies. **A**. STX-H3, young mice, Day 14 after a single IM injection **B**. STX-H3, young mice, Day 35 after two IM injections. **C**. STX-H3, young mice, IFN γ response. **D**. STX-H3, old mice, Day 14 after a single IM injection **E**. STX-H3, old mice, Day 35 after two IM injections. **F**. STX-H3, old mice, IFN γ response. N= 10/experimental group. Data are shown as mean ± SEM. **** p<0.0005, ***p<0.001, **p<0.01, *p<0.05, ns= not significant, 1-way ANOVA

### IgG ELISA for Spike

Mouse IgG antibody against SARS-CoV-2 spike was measured by an enzyme-linked immunosorbent assay (ELISA) using precoated ELISA plates (IEQ-CoV-S-RBD-IgG, RayBiotech) according to the manufacturer’s instructions, at RT. Briefly, mouse plasma samples were diluted in sample buffer (RayBiotech) and added to antigen-coated wells, in triplicates, and incubated at RT for 2 h on a shaker (200 rpm). Commercially available antibody against Spike (S1N-S58, Acro Biosystems) was used as positive controls. Plates were washed 3 times with wash buffer, incubated for 1h at RT with HRP-conjugated goat anti-mouse secondary antibodies (115-035-003, dilution 1:5000, Jackson ImmunoResearch) or anti-rabbit (111-035-003, dilution 1:5000, Jackson ImmunoResearch) diluted in assay buffer (RayBiotech). After 3 washes, plates were developed using TMB substrate (RayBiotech). After 15 min incubation, reaction was stopped by adding STOP solution and absorbance at 450 nm was recorded using a BioTeck Gen5 plate reader (Agilent). Endpoint titers were calculated as the dilution that emitted an optical density exceeding 4X the PBS control group.

### IgG ELISA for H3

Plates (Nunc MaxiSorp, Thermofisher Scientific) were coated with 10 μg/mL of H3 protein (Sino Biological, cat# 11715-V08H) in coating buffer (50 mM carbonate/ bicarbonate, pH 9.6) for 4 h at RT. Plates were rinsed, blocked with Teknova Assay Buffer (Teknova, cat # 2D5220) overnight. Plasma was diluted in Teknova Assay Buffer and incubated for 2 h at RT. After washing, plates were incubated for 1 h at RT with HRP-conjugated goat anti-mouse secondary antibodies (115-035-003, dilution 1:5000, Jackson ImmunoResearch). After 3 washes, plates were developed using TMB substrate (Surmodics/TMSK-1000-01) for 15 min, stopped with Stop Solution (acid) and absorbance values were recorded at 450 nm on a spectrophotometer (using a BioTeck Gen5 plate reader (Agilent).

## RESULTS

### Characterization of STX-H3 cell line and exosomes

293F cells were engineered to express H3 protein on their surface by lentiviral transduction. Expression of H3 on the cell surface was evaluated by flow cytometry in both adherent and suspension cells. As shown in Fig. 1A and 1B, >90% of the STX-H3 cells expressed H3. STX-H3 exosomes were purified according to the Capricor’s protocol [5, 6] and further validated. Purified STX-H3 exosomes showed an average concentration of 1.63E12 particles/mL with an expected, average diameter of 141 nm and an expected polydispersity index (PDI) of 0.164. Flow cytometry analysis showed accumulation of H3 on the exosomes (Fig. 1C), additionally confirmed by Jess Western Blot (Fig. 1D). Characterization of STX-S cells and exosomes was previously described in ([5, 6]).

### STX-H3 induces strong immunization against H3 protein in mice

The STX-H3 exosome vaccine was administered into 8-10 weeks old (study 1) and 5 mo old female mice (study 2) by intramuscular (IM) injection. Two IM injections were administered: a prime injection on day 1 followed by a second IM injection (referred to as boost injection) after a 3-week interval.

The STX-H3 vaccine induced a strong antibody response 2 weeks after the first prime injection in all animals in both studies. STX-H3 primed mice produced 15-fold (after one injection) to greater than 4500-fold (after second injection) greater H3-specific IgG than control mice (Fig. 2 A, B, D, E). No significant difference in IgG response was observed across the two groups. To characterize the T-cell response to the STX-H3 vaccine, antigen-specific Tcell responses were measured by ELISpot (Fig. 2 C, D, G, H). Splenocytes were isolated from animals at day 40 (∼3 weeks after boost (2^nd^) injection) and evaluated using ELISpot plates precoated with interferon-gamma (IFNγ). PBS was used as controls in the study. Baseline expression was compared to stimulation with 10 μg/ml of H3 protein (SinoBiological). In young mice, IFN γ response was equally increased regardless of the dose used, with a 2-to-4-fold increase in IFN γ production (Fig. 2 C). In older mice, IFN γ response was also increased at baseline in STX-H3 treated groups, with additional +2-fold increase in dose1 group (Fig. 2 F).

### STX-H3+S induces strong immunization against H3 and Spike protein in mice

We have previously shown that our STX™ Technology allows us to combine STX exosomes and formulate a “cocktail” for immunization ([6]). We combined STX-H3 and STX-S (expressing SARS-CoV-2 delta spike) exosomes and immunized mice by two IM injections: a first prime injection on day 1, followed by a booster injection on day 21 (3 wks interval).

Antibody levels against H3 and Spike were evaluated at day 14 and 35. As shown in Fig. B-E, all mice developed a strong immune-response to both antigens. Response was comparable to previous studies using single injection of either STX-H3 (Fig.2) or STX-S [5, 6]

To characterize the T-cell response to the STX-H3-S cocktail vaccine, antigen-specific T-cell responses were measured by ELISpot (Fig. 3). Splenocytes were isolated from animals at day 40 (∼3 weeks after boost (2^nd^) injection) and evaluated using ELISpot plates precoated with interleukin-4 (IL-4) or interferon-gamma (IFNγ). PBS was used as a control in the study. Baseline expression was compared to stimulation with 10 μg/ml of either H3 protein (SinoBiological) or Spike (AcroBiosystem). Both antigens were able to induce a 2-fold increase of IFN γ response, over baseline, which did not differ from PBS group (Fig. 3F-G). This data clearly suggests that our STX platform can be used for the formulation of multivalent vaccine, without antigen competition for either the antibody or T-cell response.

**Figure 3.**
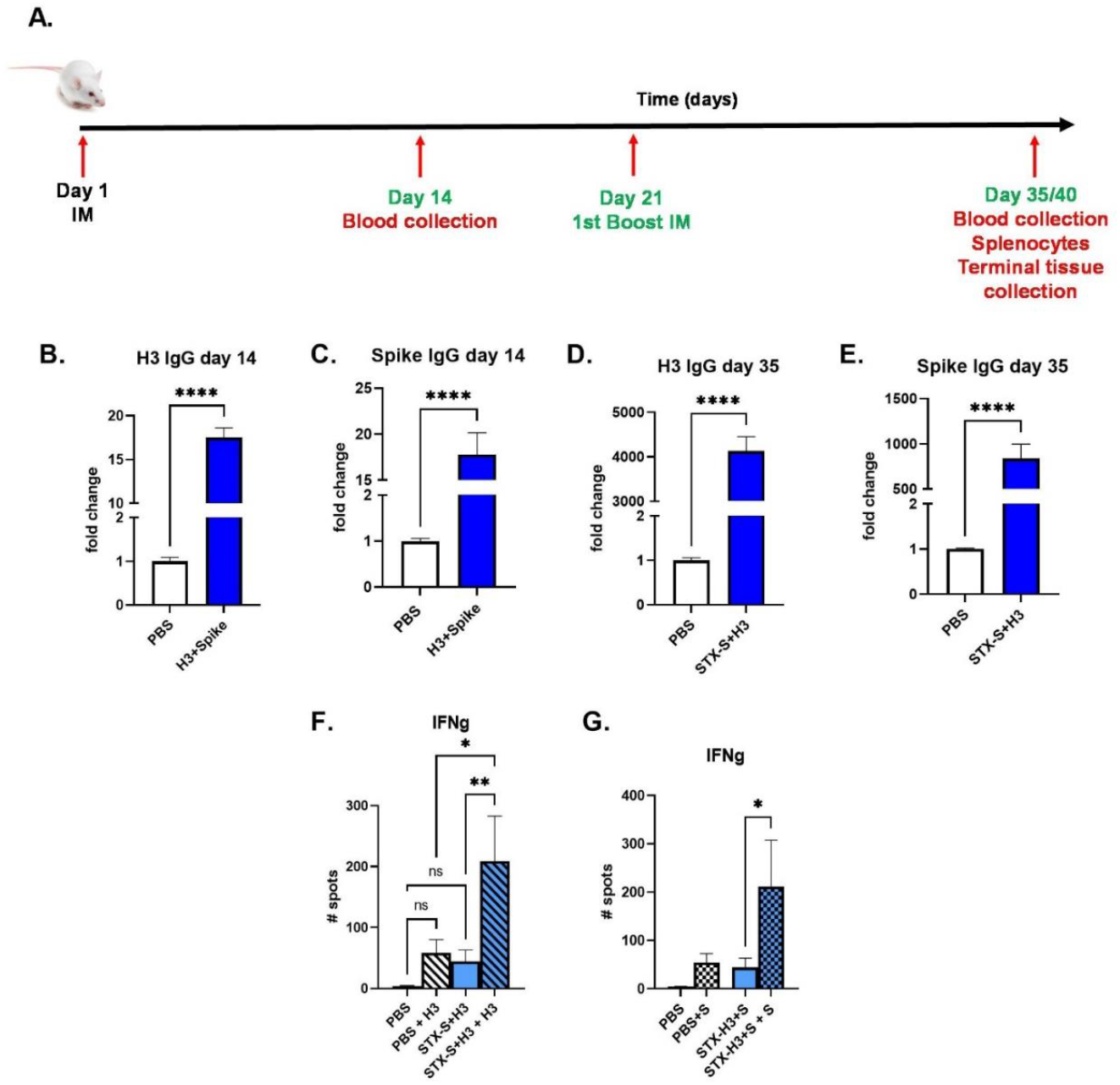
STX-H3+S exosome vaccine elicited a robust immune response. The STX-H3 vaccine induced robust expression of both anti H3 and anti-spike antibodies in mice after 1 (day 14, B, C) and 2 (day 35, D, E) IM injections as analyzed by ELISA. Additionally a strong T-cell response is observed after stimulation with either H3 or spike protein in vitro (F, G). PBS was used as a vehicle control in all studies. **A**. Schematic diagram of *in vivo* studies. **B**. Antibody against H3 on Day 14, after a single IM injection **C**. Antibody against spike on Day 14, after a single IM injection **D**. Antibody against H3 on Day 35, after a two IM injections. **E**. Antibody against spike on Day 35, after two IM injections. **F-G**. IFN γ response on day 35 after H3 (F) or spike (G) stimulation. N= 15/experimental group. Data are shown as mean ± SEM. **** p<0.0005, ***p<0.001, **p<0.01, *p<0.05, ns= not significant, 1-way ANOVA

## DISCUSSION

The constant emergence of new viruses, variants of existing viruses, and the possibilities of new pandemics have highlighted the need for multivalent vaccines. These multivalent vaccines must be able to strongly protect the population not only against the rapid spread of the viruses but also against their consequences on human health and society as evidenced by the impact of the COVID-19 pandemic.

The innate characteristics of exosomes can enable them to be the next generation vaccine: they show high bioavailability, remarkable biocompatibility, and low immunogenicity, making exosomes a very promising drug delivery candidate [7]. Because of their lipid bilayer, exosomes could potentially cross any biological barriers and deliver their cargo, endogenously or exogenously expressed [8]. Exosomes showed a high stability at physiological pH and temperature, a unique advantage as delivery system for vaccinology. Moreover, because exosomes are a natural product of any cell type, they are less likely to induce an adverse response, improving their safety profile.

We have previously showed that our Stealth-X™ platform can easily and rapidly provide a multi-protein-based vaccine delivered by exosomes because of its easy scale-up manufacturing process, and can produce a strong antibody response, demonstrated by neutralizing antibodies and a strong T-cell response with a minimum number of injections [5, 6]. The platform allows multiplexing: because of the small amount of protein needed to induce an immune response, exosomes carrying different antigens can be individually prepared and subsequently combined according to the desired dose.

Here we apply the STX technology to produce a multivalent protein-based exosome delivered vaccine, against both Influenza and SARS-CoV-2. Previously, we demonstrated that STX-S and STX-N can elicit robust humoral and cellular immune responses against different variants of SARS-CoV-2 viruses[5, 6]. Similarly in the present study, monovalent STX-H3 vaccination induced both IgG and T-cell responses against H3. To test the potential benefit of the bivalent vaccine, hemagglutinin H3 and SARS-CoV-2 Spike proteins were individually engineered on the exosome surface and mixed before injection. This ‘cocktail’ vaccine was able to induce a strong immune-response in mice, with both antibody and T cell response, without detectable immune-competition between antigens. These represent two important characteristics of a potent vaccine, where administration can activate both the innate and cellular immune response, for a long lasting and effective protection against infections. Furthermore, the additive immunogenicity induced by the administration of the bivalent STX-S and STX-H3 vaccine demonstrated the great potential of the StealthX™ vaccine technology in controlling a broad range of infectious diseases in a single vaccine formulation.

Our data suggest that exosomes are ideal vehicles for vaccination because they can safely deliver the antigen of interest (exogenous protein) efficiently by mimicking the natural viral infection.

Vaccination is a cost-effective public health measure which can prevent disease spread and reduce disease burden. Despite the lifesaving benefits of increased vaccine uptake, vaccine hesitancy and skepticism are still longstanding barriers to the health of individuals (<u>Centers for Disease Control and Prevention, 2019</u>). The nanogram dose used in our formulations allows the multiplexing of several antigens: this could reduce the number of injections, with important cost-benefits, and possibly improving the vaccination rate with a broader population immunization and significant impact on health.

In conclusion, we rapidly generated a dual-antigen vaccine with efficacy against multiple SARS-CoV-2 and Influenza, that has broader immune capability, and could be used as a booster vaccine to the existing immunity generated by previously approved vaccines. Our StealthX vaccine technology, which uses rapidly engineered and manufactured exosomes to deliver nanograms of single or multiplexed viral antigens to elicit a strong, broad immune response without any adjuvants or lipid nanoparticles, has the ability to revolutionize the next generation of vaccines.

## Author contribution statements

MC contributed to the design, implementation, and analysis of all research in this paper and to the writing of the manuscript. MS contributed to all the design of STX constructs; LH and YL produced the cell lines, carried the in vitro experiments. MJL, MY, JA, CL, CJ produced the exosomes used in the study. KE, MS contributed to the design and review of studies and the writing of the manuscript. MC, LH, JL, JBN, CM, MJL, JA, MY, CL, CJ, RT carried out the experiments.

